# Enhancing mitochondrial activity in neurons protects against neurodegeneration in CNS inflammation

**DOI:** 10.1101/2020.06.19.161091

**Authors:** Sina C. Rosenkranz, Artem A. Shaposhnykov, Simone Träger, Jan Broder Engler, Maarten E. Witte, Vanessa Roth, Vanessa Vieira, Nanne Paauw, Simone Bauer, Celina Schwencke-Westphal, Lukas Bal, Benjamin Schattling, Ole Pless, Jack van Horssen, Marc Freichel, Manuel A. Friese

**Author notes:** **Corresponding author:** Manuel A. Friese, Institute of Neuroimmunology and Multiple Sclerosis (INIMS), University Medical Center Hamburg-Eppendorf, Falkenried 94, 20251 Hamburg, Germany.

## Abstract

Central nervous system (CNS) inflammation in multiple sclerosis (MS) drives neuro-axonal loss resulting in irreversible disability. While transcripts of mitochondrial genes are strongly suppressed in neurons during CNS inflammation, it is unknown whether this results in mitochondrial dysfunction and whether interventions that increase mitochondrial function can rescue neurodegeneration. Here we show that suppression of mitochondrial gene transcripts in inflamed neurons was predominantly affecting genes of the electron transport chain resulting in impaired mitochondrial complex IV activity. This was associated with posttranslational inactivation of the transcriptional co-regulator peroxisome proliferator-activated receptor gamma co-activator 1-α (PGC-1α). Neuronal overexpression of *Pgc-1α* led to increased numbers of mitochondria, complex IV activity and elevated maximum respiratory capacity. Moreover, *Pgc-1α-* overexpressing neurons showed a higher mitochondrial membrane potential that related to an improved calcium buffering capacity. Accordingly, neuronal deletion of *Pgc-1α* aggravated neurodegeneration during experimental autoimmune encephalomyelitis (EAE), while neuronal overexpression of *Pgc-1α* ameliorated EAE disease course and preserved neurons. Our study provides systemic insights into mitochondrial dysfunction in neurons during inflammation and commends elevation of mitochondrial activity as a promising neuroprotective strategy.

## INTRODUCTION

Multiple sclerosis (MS) is a chronic inflammatory disease of the central nervous system (CNS) and the most frequent non-traumatic cause of neurological impairment during early adulthood. Neuronal loss occurs already from disease onset and correlates best with irreversible disability in MS patients (Fisher *et al*, 2008; Tallantyre *et al*, 2010; Fisniku *et al*, 2008). While there has been substantial progress in the understanding and treatment of the immune response, the pathogenesis of concurrent neuronal damage is incompletely understood. Recently two drugs, ocrelizumab and siponimod, have been licensed for the treatment of progressive MS patients, however, their mode of action relies on immune regulation (Kappos *et al*, 2018; Montalban *et al*, 2016). Since there are currently no therapies available to counteract the progression of neurodegeneration in MS patients (Feinstein *et al*, 2015; Dendrou *et al*, 2015) and repurposing of drugs to directly target neurodegeneration in MS has been disappointing (Chataway *et al*, 2020), a better understanding of the molecular processes that determine neuronal cell loss in MS are urgently needed.

Neuronal loss in MS and its animal model, experimental autoimmune encephalomyelitis (EAE), has been associated with enhanced production of reactive oxygen and nitrogen species by immune cells and with increased iron accumulation in the gray matter (Stephenson *et al*, 2014). Both processes can lead to damage of neuronal mitochondria with subsequent metabolic failure (Campbell *et al*, 2011). Moreover, disruption of neuronal ion homeostasis (Craner *et al*, 2004; Friese *et al*, 2014) and aggregation of neuronal proteins (Schattling *et al*, 2019) consume high amounts of energy that might further drive neuroaxonal injury. Furthermore, excessive activation of calcium-dependent processes and neuronal calcium overload seems to be another important component of neuronal injury (Friese *et al*, 2007; Schattling *et al*, 2012; Witte *et al*, 2019). While identifying druggable targets that specifically induce neuronal resilience has been extremely difficult due to insufficient insights into key modulators, mitochondria could serve as an important hub, as they are pivotal for both energy production and calcium homeostasis (Vafai & Mootha, 2012).

Mitochondria usually have a high calcium buffering capacity, which is driven by their negative membrane potential, which is generated by the activity of oxidative phosphorylation (Kann & Kovács, 2007; Zorova *et al*, 2018). However, overload of mitochondria with calcium as a consequence of CNS inflammation can result in inappropriate activation of the mitochondrial permeability transition pore (PTP) with subsequent mitochondrial swelling and cell death (Rizzuto *et al*, 2012; Giorgi *et al*, 2018), which is one of the neuropathological hallmarks in neurons during EAE (Nikić *et al*, 2011). Inhibiting the mitochondrial matrix protein cyclophilin D (CyPD), a regulator of the PTP can partially counteract this dysregulation (Forte *et al*, 2007). Additionally, to counteract ion imbalance, a higher activity of ATP-dependent ion pumps would be required. However, post mortem studies of MS patients’ CNS tissue revealed a compromised neuronal ATP production by oxidative phosphorylation, as decreased mitochondria complex IV activity in demyelinated axons and neurons were detected (Mahad *et al*, 2009; Campbell *et al*, 2011). Similarly, mitochondrial gene expression is suppressed in motor neurons of the spinal cord in EAE mice (Schattling *et al*, 2019), further supporting a comprised mitochondrial function during CNS inflammation.

Thus, excess neuroaxonal calcium together with mitochondrial dysfunction has been postulated to trigger neuroaxonal injury observed in MS and EAE. This would be predicted to lead to elevated levels of calcium within neuronal and axonal mitochondria further perpetuating mitochondrial dysfunction and neuronal injury. However, it is currently unknown whether interventions that increase mitochondrial energy production in neurons can rescue neurodegeneration during CNS inflammation.

Here we discovered by an unsupervised survey of neuronal gene expression during CNS inflammation that genes involved in the electron transport chain, especially in complex I and IV are repressed in motor neurons. While this is not accompanied by a decrease of mitochondrion numbers in motor neuronal somata, we detected a decrease in neuronal mitochondrial complex IV activity. This implies a lower activity of neuronal mitochondria during EAE and we were able to associate that with a posttranslational inactivation of peroxisome proliferator-activated receptor gamma coactivator 1-alpha (PGC-1α), one of the master regulators of mitochondrial numbers and function. Notably, neuronal overexpression of *Pgc-1α* led to an increase of mitochondrial activity, especially in complex IV activity and substantially elevated calcium buffering capacity. Subjecting these mice to EAE resulted in a significantly better recovery from clinical disability in comparison to wild-type controls. Together, induction of neuronal mitochondrial activity commends as a promising therapeutic approach to counteract inflammation-induced neurodegeneration.

## RESULTS

### Impaired neuronal oxidative phosphorylation in inflamed spinal cords

Since we have previously discovered that mitochondrial gene transcripts are markedly suppressed in motor neurons during CNS inflammation in the EAE model (Schattling *et al*, 2019), we first investigated whether unifying biological processes are dysregulated. Therefore, we derived a gene expression signature of motor neurons in spinal cord during CNS inflammation (Schattling *et al*, 2019) and tested for gene set enrichment of biological process gene ontology (GO) terms. While the majority of terms was enriched during inflammation, only a confined group of 29 terms showed strong de-enrichment (Fig 1A, dashed line) and was dominated by terms representing mitochondrial function (Fig 1B). A recurring theme was the suppression of the electron transport chain (ETC; Fig 1C). Suppression of gene transcripts that are involved in the ETC, which is the key mechanism of oxidative phosphorylation, can either result in reduced number of mitochondria or an impairment of mitochondrial function. Therefore, we quantified mitochondria content in the soma of motor neurons in EAE mice at the chronic stage in comparison to healthy control mice, but did not detect any difference in mitochondria area normalized to neuronal size (Fig 1D, Fig EV1A). However, we detected a tendency towards an increase in overall numbers of motor neuronal mitochondria (Fig EV1B) that was based on an increased size of the neuronal cell body during EAE (Fig 1E). By contrast, the size of mitochondria did not differ in motor neurons of healthy and EAE mice (Fig EV1C).

**Figure 1:**
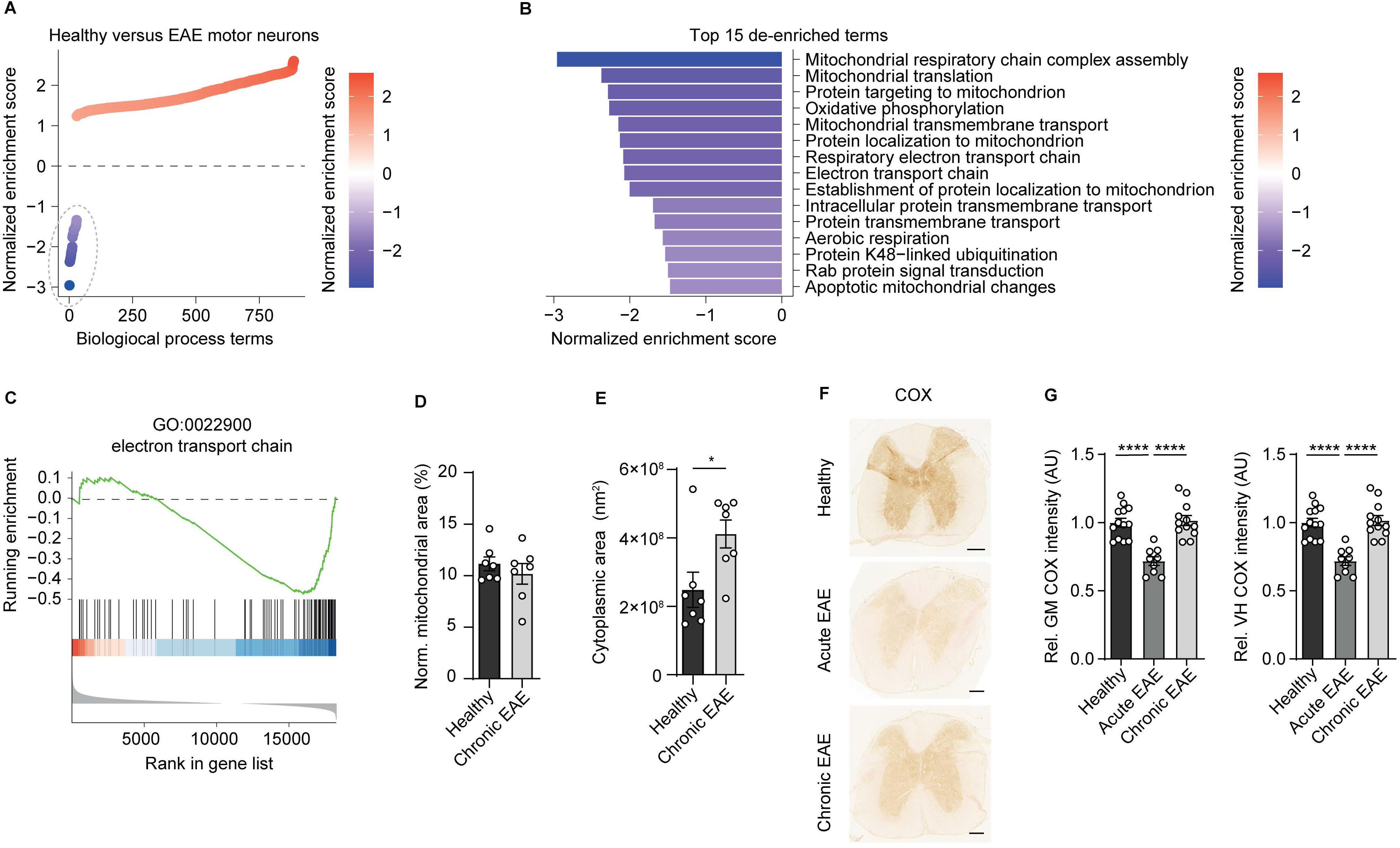
Reduced activity of oxidative phosphorylation in spinal cord neurons during EAE. (**A**) Gene set enrichment analysis (GSEA) of biological process gene ontology (GO) terms in motor neurons during CNS inflammation in the EAE model. Dashed line indicates 29 de-enriched terms. (**B**) Top 15 de-enriched GO terms with normalized enrichment score. (**C**) GSEA plot of the de-enriched term ‘GO:0022900 electron transport chain’. (**D**) Transmission electron microscopy (TEM) analysis of motor neuronal mitochondrial content normalized to size of healthy (n = 3) and chronic EAE (n = 3) mice (2–3 cells per mice). Bars show mean values ± s.e.m. (**E**) Transmission electron microscopy (TEM) analysis of motor neuronal size of healthy (n = 3) and chronic EAE (n = 3) mice (2–3 cells per mice). Bars show mean values ± s.e.m. (**F**) Representative images of COX histochemistry of cervical spinal cord sections of healthy, acute and chronic EAE mice. Scale bar: 250 μm. (**G**) Quantification of COX histochemistry of cervical spinal cord gray matter (GM) and ventral horn (VH) of healthy (n = 5), acute (n = 3) and chronic (n = 5) EAE mice (2–3 stainings per mice) normalized to neuronal nuclei (NeuN)-positive neurons. Bars show mean values ± s.e.m. Statistical analysis in (**D**) and (**E**) was performed by unpaired, two-tailed Student’s t-test, in (**G**) by one-way ANOVA following Tukey’s post-hoc test for multiple comparisons. *P < 0.05, ****P < 0.0001.

Dismissing differences in mitochondrial content, we reasoned that downregulation of ETC transcripts could result in a compromised oxidative phosphorylation activity. By analyzing COX histochemistry which represents complex IV activity, we detected a significant decrease in absolute neuronal complex IV activity in the entire gray matter and ventral horn of acute EAE animals (day 15 post immunization) and in chronic EAE animals (day 30 post immunization) in comparison to healthy control mice (Fig 1F). Normalization to NeuN-positive neurons revealed that the reduced complex IV activity at day 30 was due to neuronal loss, whereas at day 15 the reduced complex IV activity was independent of neuronal numbers (Fig 1G).

### Inactivation of PGC-1α contributes to mitochondrial dysfunction in CNS inflammation

Next we interrogated whether a unifying mechanism can explain the inflammation-induced downregulation of gene transcripts that are involved in ETC with consecutively impaired complex IV activity in neurons at the acute stage of EAE. Notably, we detected that many of the genes which drive the downregulation of the ETC theme are usually induced by the transcriptional coactivator *Pgc-1α* (Puigserver *et al*, 1998; Lucas *et al*, 2014) (Fig 2A). Consequently, we asked whether *Pgc-1α* could be the sought-after unifying factor that is functionally disturbed. Motor neuronal *Pgc-1α* expression itself was not altered in our previously published data set comparing acute EAE to controls (Fig 2B). After confirming this result by an additional translating ribosome affinity purification (TRAP) of motor neuronal transcripts followed by qPCR (Fig 2C) and immunoblot of PGC-1α of the cervical spinal cord of EAE and healthy mice (Fig 2D, E), we concluded that transcriptional or translational *Pgc-1α* /PGC-1α regulation cannot be the explanation of the observed ETC downregulation.

**Figure 2:**
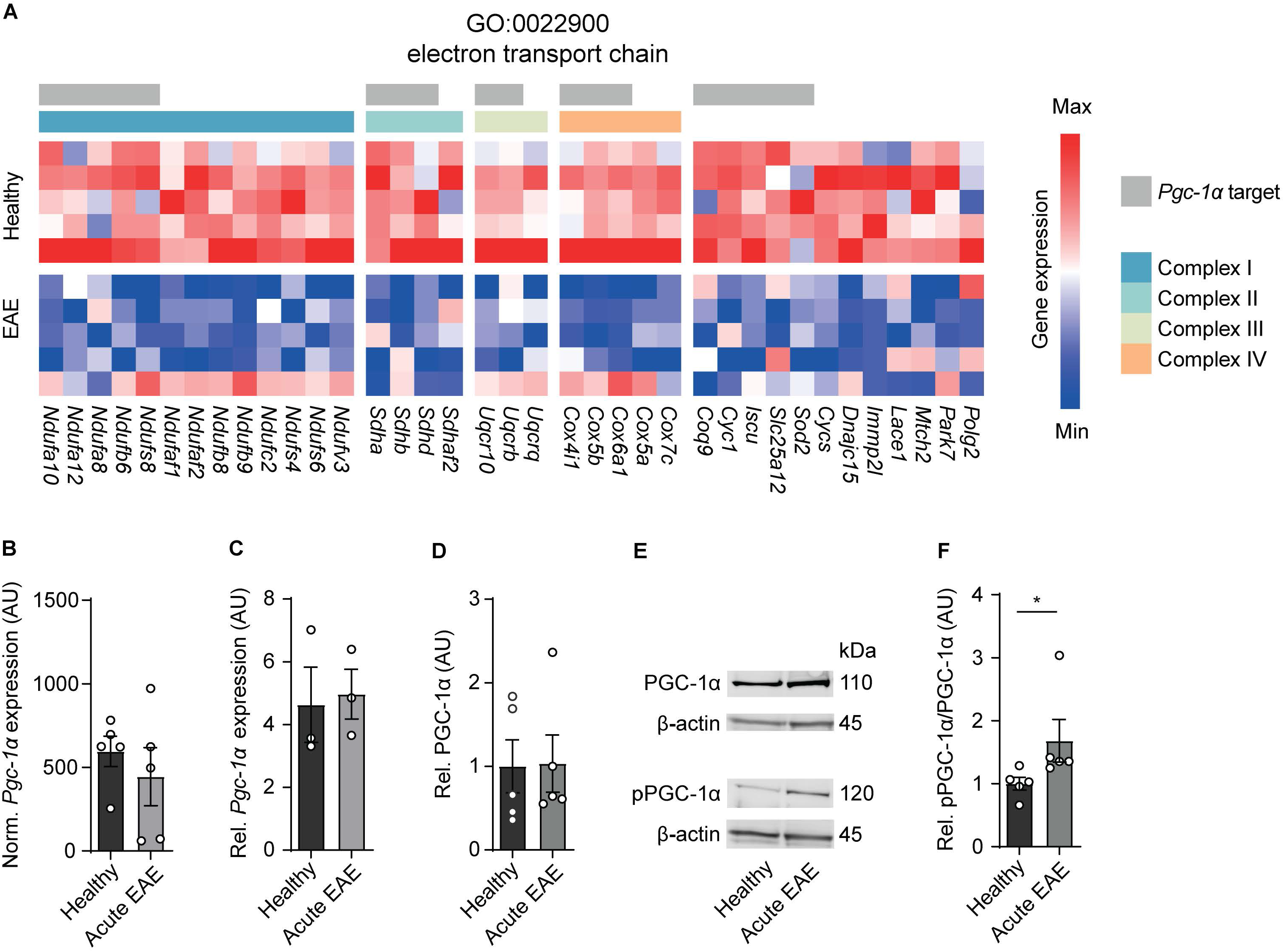
Inactivation of PGC-1α in spinal cord neurons during EAE. (**A**) Heatmap of genes driving the de-enrichment of ‘GO:0022900 electron transport chain’. Heatmap is annotated for electron transport chain complexes and target genes induced by the transcription factor *Pgc-1α* (Lucas *et al*, 2014). (**B**) Normalized RNA-seq expression of *Pgc-1α* in healthy (n = 5) and acute EAE (n = 5) motor neurons. Bars show mean values ± s.e.m. (**C**) Relative qPCR mRNA expression of *Pgc-1α* to *Tbp* in healthy (n = 3) and acute EAE (n = 3) motor neurons. Bars show mean values ± s.e.m. (**D**) Quantification of PGC-1α protein in cervical spinal cords of healthy (n = 5) and acute (n = 5) EAE mice. Each sample was normalized to its β-actin. Bars show mean values ± s.e.m. (**E**) Representative immunoblots of PGC-1α, phosphorylated PGC-1αS570 (pPGC-1α) and corresponding β-actin of cervical spinal cords of healthy and acute EAE mice. (**F**) Quantification of phosphorylated PGC-1αS570 (pPGC-1α) in relation to PGC-1α total protein in cervical spinal cords of healthy (n = 5) and acute (n = 5) EAE mice. Each sample was normalized to its β-actin. Bars show mean values ± s.e.m. Statistical analysis in (**B**) and (**C**) was performed by unpaired, two-tailed Student’s t-test, in (**D**) and (**F**) by unpaired, two-tailed Mann-Whitney test; *P < 0.05.

Besides quantity, PGC-1α can be post-translationally inactivated by phosphorylation at serine 570 (Li *et al*, 2007; Xiong *et al*, 2010) and thereby preventing the recruitment of PGC-1α to its regulated promoters, which we tested next. After validation of phosphorylation specificity of the pPGC-1αS570 antibody (Fig EV2A), we detected a significant increase in phosphorylated-serine570 PGC-1α in spinal cords of acute EAE mice in comparison to healthy control mice (Fig 2E, F). Thus, we concluded that due to the phosphorylation of PGC-1α during acute stage of EAE which is associated with downregulation of its target gene transcripts and compromised mitochondrial function, *Pgc-1α* could be a potential target to rescue mitochondrial function in EAE affected motor neurons.

### Neuronal Pgc-1α induction augments mitochondrial function

Next we reasoned that elevating neuronal *Pgc-1α* could restore mitochondrial function and serve as a strategy to improve neuronal resilience in CNS inflammation. We utilized transgenic mice with a neuron-specific overexpression of *Pgc-1α* driven under the Thy1 promoter (*Pgc-1α^T^*^/T^) (Mudò *et al*, 2012), which is expressed in mature neurons (Fig EV3A). We made sure that elevated transgenic DNA copies of Pgc-1α in neurons (Fig EV3B) resulted in elevated mRNA expression of *Pgc-1α* in cortex, hippocampus and spinal cord as well as in primary neurons of *Pgc-1α*^T/T^ mice in comparison to wild-type mice (Fig EV3C, D). Additionally, we revealed that *Pgc-1α*^T/T^ mice showed an increase of *Pgc-1α*-dependent transcriptional targets in primary neurons (Fig EV3E, F) and *cytochrome c oxidase subunit 4* (*Cox4i1*) and *citrate synthase* (*Cs*) in CNS tissue (Fig EV3G, H, I). Increased mRNA level led to a corresponding increase in protein amount of PGC-1α in primary neurons (Fig EV3J) and spinal cord, which we confirmed by staining for the transgenically fused FLAG-tag (Fig EV3K). Accordingly, neuronal *Pgc-1α* overexpression resulted in increased mitochondrial content in CNS tissue (Fig 3A) and primary neurons (Fig 3B).

**Figure 3:**
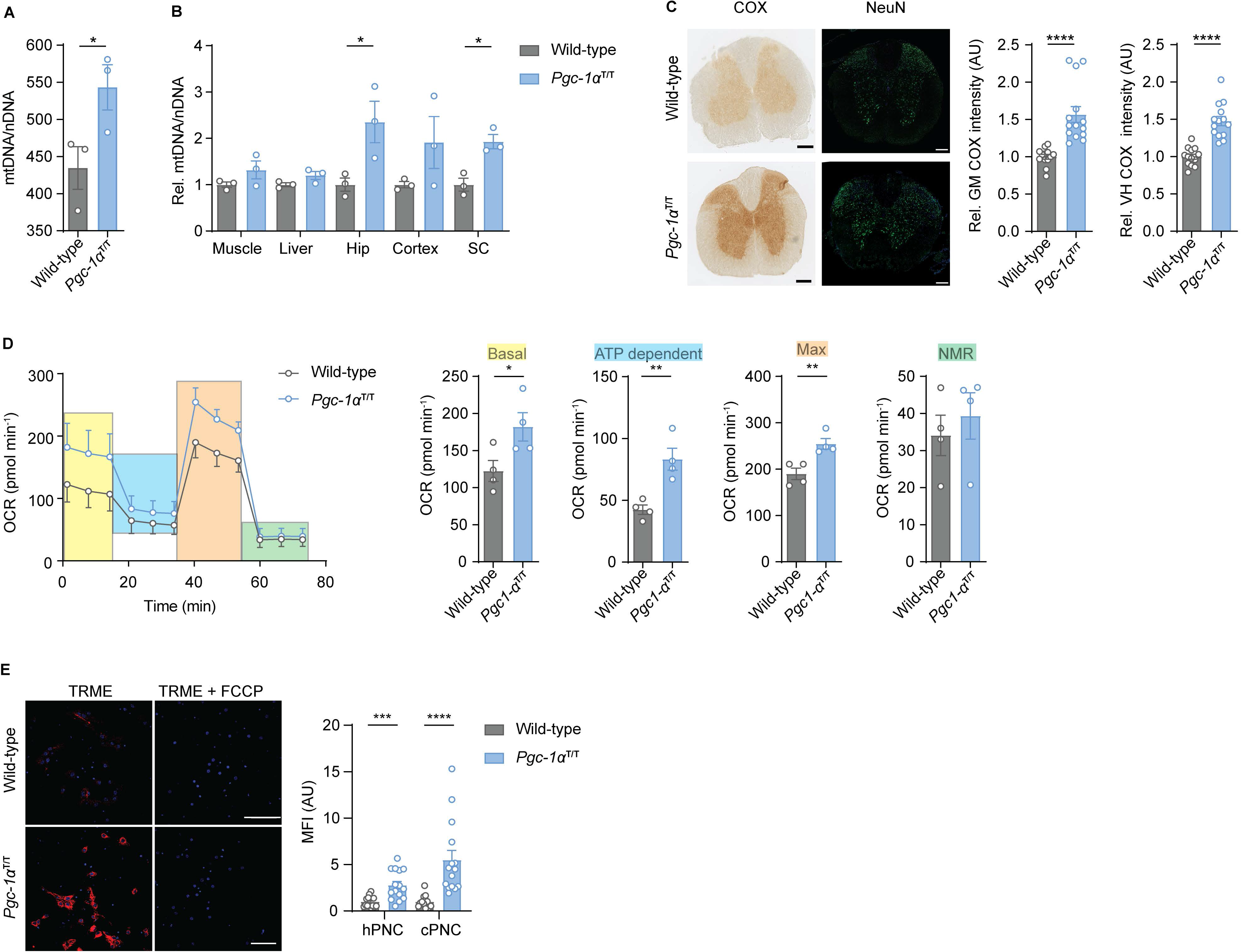
Neuronal overexpression of *Pgc-1α* increases neuronal mitochondrial activity. (**A**) Mitochondrial DNA copy numbers (mtDNA) relative to diploid nuclear chromosomal DNA (nDNA) in hippocampal primary neuronal cultures (hPNC) (DIV 14) of wild-type (n = 3) and *Pgc-1α*^T/T^ (n = 3) mice. Bars show mean values ± s.e.m. (**B**) Mitochondrial DNA copy numbers (mtDNA) relative to diploid nuclear chromosomal DNA (nDNA) in muscle, liver, hippocampus (Hip), cortex and cervical spinal cord (SC) of wild-type (n = 3) and *Pgc-1α*^T/T^ (n = 3) mice. Bars show mean values ± s.e.m. (**C**) Representative images and quantification of COX histochemistry of cervical spinal cord gray matter (GM) or ventral horn (VH) of wild-type (n = 5) and *Pgc-1α*^T/T^ (n = 6) mice (2–3 stainings per mice) normalized to neuronal nuclei (NeuN)-positive neurons. Bars show mean values ± s.e.m. Scale bar: 250 μm. (**D**) Profile and quantification of oxygen consumption rate in hippocampal primary neuronal culture (hPNC) (DIV 14) of wild-type (n = 4) and *Pgc-1α*^T/T^ (n = 4) mice. Yellow: Basal respiration (Basal); blue: ATP-dependent respiration (ATP dependent); orange: Maximal respiratory capacity (Max); green: Non-mitochondrial respiration (NMR). Bars show mean values ± s.e.m. (**E**) Representative images and mean fluorescence intensity (MFI) quantification of TMRE mitochondrial membrane potential assay of hippocampal primary neuronal culture (hPNC) (DIV 14) of wild-type (n = 3) and *Pgc-1α*^T/T^ (n = 3) mice (5 cells per culture). FCCP was used as ionophore uncoupler of oxidative phosphorylation. Bars show mean values ± s.e.m. Scale bar: 10 μm. Statistical analysis in (**A**-**E**) was performed by unpaired, two-tailed Student’s t-test. *P < 0.05. **P < 0.01, ***P < 0.001, ****P < 0.0001.

To determine the mitochondrial activity in these mice we analyzed whether *Pgc-1α*-mediated *Cox4i1* induction resulted in increased complex IV activity. We detected an elevated complex IV activity in the gray matter and ventral horn of *Pgc-1α*^T/T^ mice, which was not due to a higher number of NeuN-positive neurons (Fig 3C). Metabolically, primary neurons of *Pgc-1α*^T/T^ mice showed an elevated oxygen consumption rate and maximum respiratory capacity (Fig 3D), which was independent of neuronal numbers (Fig EV3L). In concordance with higher activity of oxidative phosphorylation, primary neurons of *Pgc-1α*^T/T^ mice retained higher TMRE fluorescence intensity than wild-type controls, representing a hyperpolarized mitochondrial membrane potential (Fig 3E).

### Induction of neuronal Pgc-1α improves neuronal calcium buffering capacity

As CNS inflammation results in substantial neuronal calcium influx (Witte *et al*, 2019) leading to neurodegeneration, which could potentially be buffered by mitochondria, we next asked whether increased neuronal mitochondrial activity and membrane potential in *Pgc-1α*^T/T^ neurons could alleviate toxic calcium levels. By analyzing spontaneously active cortical primary neuronal cultures (cPNC) transduced with the virally encoded calcium indicator GCaMP6f (Fig 4A) we recorded a significantly faster clearance of intracellular calcium concentrations and decreased amount of mean calcium in spontaneously active *Pgc-1α*^T/T^ primary neurons in comparison to wild-type control neurons, whereas the amplitude and number of calcium transients did not differ (Fig 4B, C). The results were confirmed with a chemical calcium indicator (Fig 4D and Appendix Video 1).

**Figure 4:**
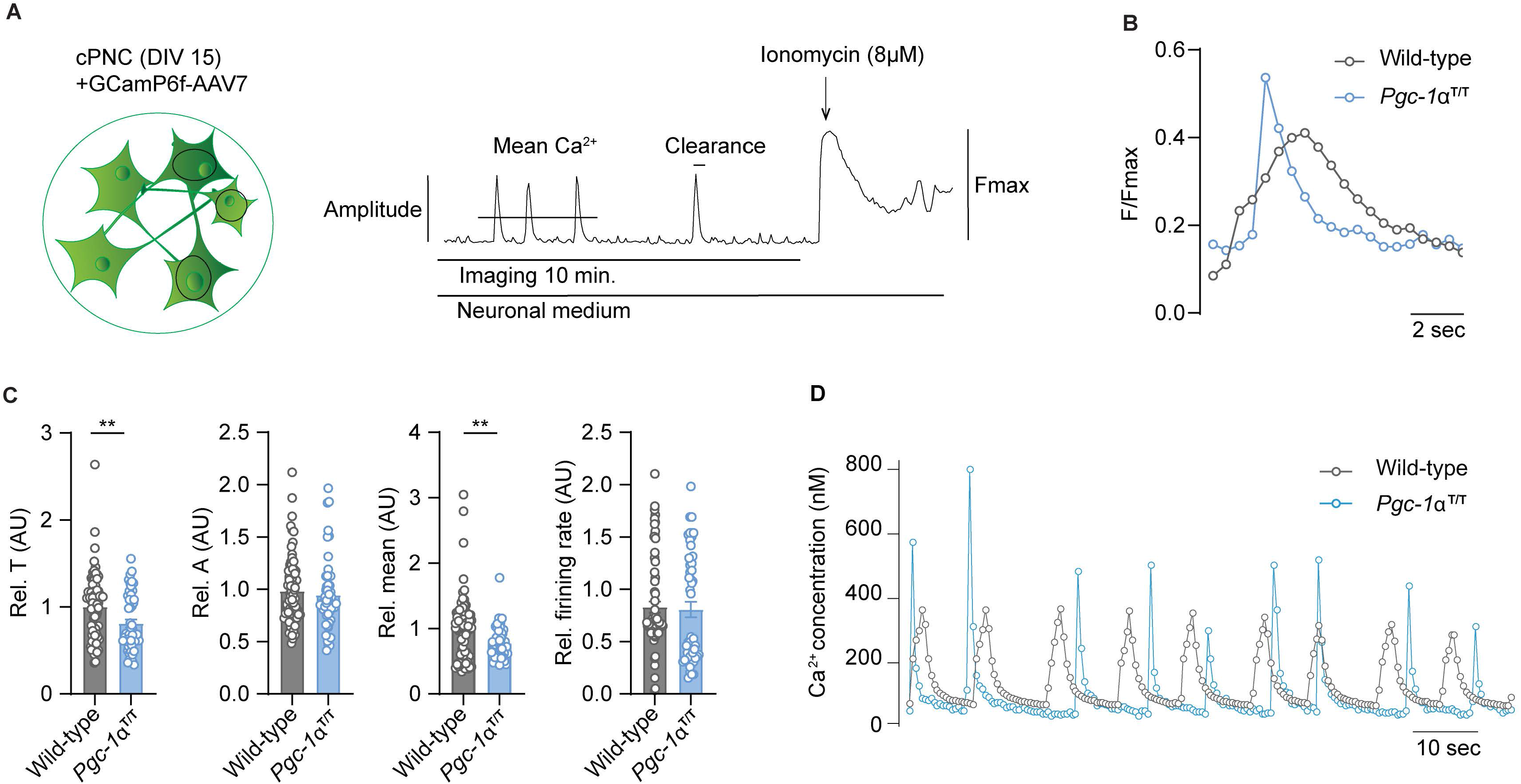
Neuronal overexpression of *Pgc-1α* improves neuronal calcium buffering. (**A**) Experimental approach for analysis of Ca^2+^ signaling. GCamp6f fluorescence was recorded for 4870 seconds in spontaneously active cortical primary neuronal culture (cPNC) (DIV15) prior to ionomycin application used for signal normalization. (**B**) Representative cytosolic calcium spikes of GCaMP6f-transduced spontaneously active cortical primary neuronal culture (cPNC) (DIV15) of wild-type and *Pgc-1α*^T/T^ mice normalized to fluorescence of calcium-saturated indicator (Fmax). (**C**) Quantification of calcium transient decay constant Tau (T) as an indicator of cytosolic calcium clearance time, of calcium transient amplitude (A), and of mean cytosolic calcium signal intensity and of the number of cytosolic calcium transients presented as firing rate of GCaMP6f-transduced cortical primary neuronal culture (cPNC) (DIV 15) of wild-type (n = 74 cells from 3 different mice) and *Pgc-1α*^T/T^ (n = 51 cells from 3 different mice). Bars show mean values ± s.e.m. (**D**) Representative cytosolic calcium spikes of Fluo-4-stained cortical primary neuronal cultures (cPNC) (DIV14–16) of wild-type and *Pgc-1α*^T/T^ mice. Statistical analysis in (**C**) was performed by unpaired, two-tailed Student’s t-test. **P < 0.01, ****P < 0.0001.

### Elevated neuronal mitochondrial activity ameliorates neurodegeneration in CNS inflammation

As we assumed that inactivation of neuronal *Pgc-1α* during CNS inflammation aggravates mitochondrial dysfunction and spurs neuronal vulnerability, we first tested whether this hypothesis holds true in *Pgc-1α*-deficient neurons. We used *Pgc-1α*^flx/flx^ × Eno2^Cre+^ mice to specifically delete *Pgc-1α* in neurons as Eno2 is widely expressed in neurons (Fig EV4A). Although *Eno2* expression is also present by immune cells (Heng *et al*, 2008), we could exclude *Pgc-1α* expression in immune cells (Fig EV4B, Heng et al., 2008) therefore securing that *Pgc-1α*^flx/flx^ × Eno2^Cre+^ mice will not result in altered immune responses. Neuronal deletion of *Pgc-1α* resulted in decreased *Pgc-1α* mRNA levels (Fig EV4C) and Pgc-1α-regulated downstream targets in hippocampus (Fig EV4D), cortex (Fig EV4E) and spinal cord (Fig EV4F). Consequently, neuron-specific *Pgc-1α* knockout mice (*Pgc-1α*^flx/flx^ × Eno2^Cre+^) showed a more severe EAE disease course (Fig 5A) and increased neuronal loss in the gray matter and ventral horn in comparison to *Pgc-1α*^flx/flx^ control EAE mice (Fig 5B).

**Figure 5:**
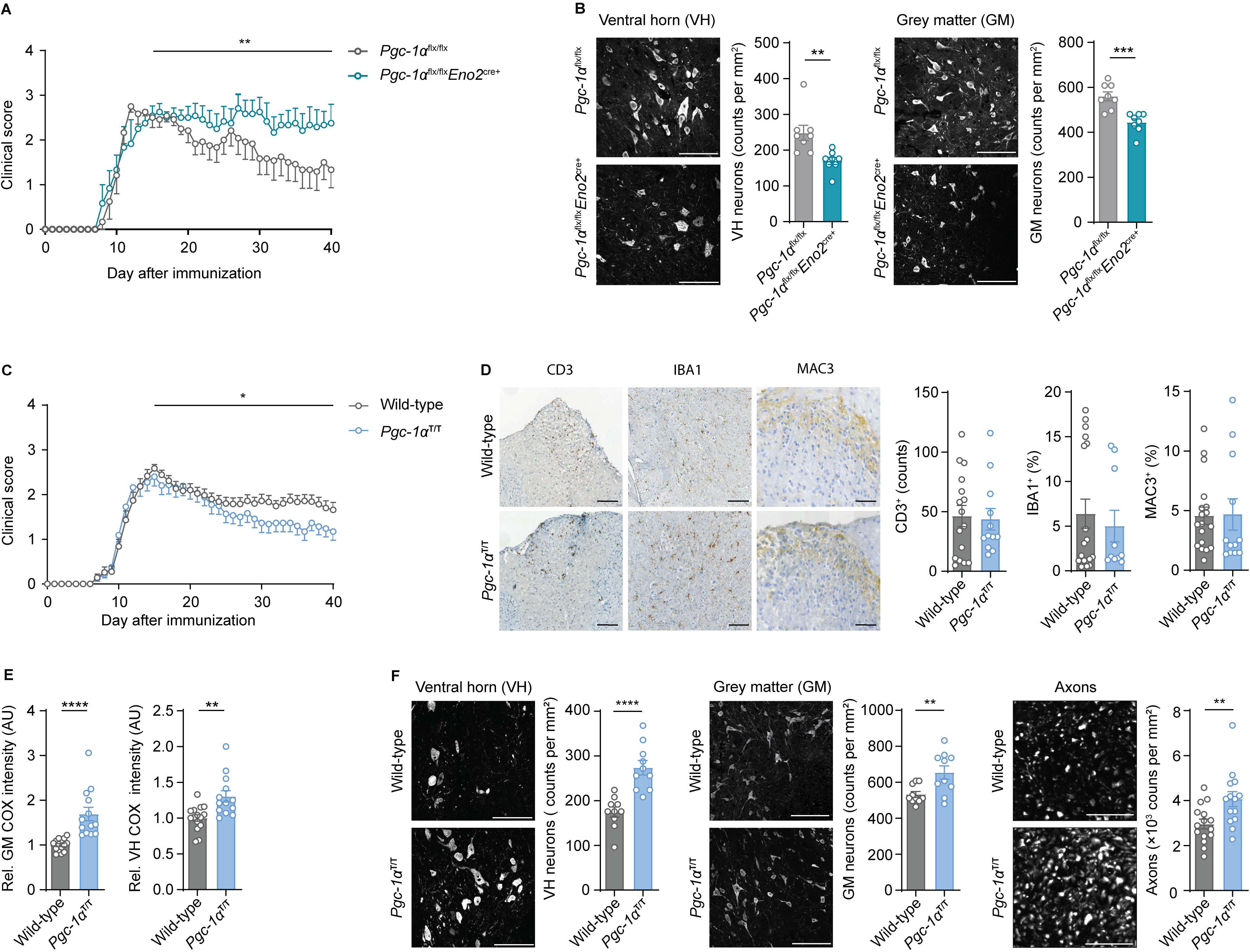
Neuronal *Pgc-1α* levels determine neuronal injury and clinical disability in EAE. (**A**) Mean clinical scores of *Pgc-1α*^flx/flx^ (n = 6) and *Pgc-1α*^flx/flx^ Eno2^cre+^ (n = 6) mice undergoing EAE. Curves show mean ± s.e.m. (**B**) Immunohistochemical stainings of surviving NeuN-positive neurons in cervical spinal cord gray matter (GM) or ventral horn (VH) of *Pgc-1α*^flx/flx^ (n = 4) and *Pgc-1α*^flx/flx^ Eno2^cre+^ (n = 4) mice (2 areas per mice) at day 40 post immunization with quantification. Bars show mean values ± s.e.m. Scale bar: 100 μm. (**C**) Mean clinical scores of wild-types (n = 31) and *Pgc-1α*^T/T^ (n = 27) mice undergoing EAE. Curves show mean ± s.e.m., all pooled from three independent experiments. (**D**) Histopathological stainings of T cells (CD3), microglia (IBA1) and macrophages (MAC-3) in cervical spinal cord sections of wild-type (n = 6) and *Pgc-1α*^T/T^ (n = 5) mice (2–3 stainings per mice) at day 15 post immunization with quantifications. Bars show mean values ± s.e.m. Scale bar: 250 μm and 100 μm. (**E**) Quantification of COX histochemistry of cervical spinal cord gray matter (GM) or ventral horn (VH) of wild-type (n = 6) and *Pgc-1α*^T/T^ (n = 4) mice (2–3 stainings per mice) at acute stage of EAE normalized to neuronal nuclei (NeuN)-positive neurons. Bars show mean values ± s.e.m. Scale bar: 250 μm. (**F**) Representative images of immunohistochemical stainings of surviving NeuN-positive neurons in cervical spinal cord gray matter (GM) or ventral horn (VH) and surviving neurofilament-positive axons in the dorsal columns of wild-type (n = 5) and *Pgc-1α*^T/T^ (n = 5) (2 areas per mice for NeuN, 3 areas per mice for neurofilament) at day 40 post immunization with quantification. Bars show mean values ± s.e.m. Scale bar: 100 μm and 50 μm. Statistical analysis in (**A**) and (**C**) was performed by one-tailed Mann-Whitney U test of area under the curve (AUC) starting at peak (day 15) of disease; in (**B**), (**D**), (**E**) and (**F**) by unpaired, two-tailed Student’s t-test. *P < 0.05; **P < 0.01; ***P < 0.001; ****P < 0.0001.

As we observed that neuronal *Pgc-1α* overexpression overruled inflammation-associated inactivation with increased oxidative phosphorylation and improved calcium buffering capacities, we next tested its translatability to the preclinical MS model. EAE induction in *Pgc-1α*^T/T^ and wild-type littermate control mice resulted in a comparable maximum score in the acute phase of the disease. However, *Pgc-1α*^T/T^ mice showed a significantly better recovery from clinical disability in comparison to wild-type controls (Fig 5C). Notably, this protection is unlikely to be driven by an impaired immune response, as numbers of infiltrating CD3^+^ T cells, activated IBA1^+^ microglia and MAC-3^+^ macrophages were similar to WT mice (Fig 5D). Moreover, neuronal overexpression of *Pgc-1α* resulted in a rescue of complex IV activity during acute EAE in the entire gray matter and ventral horn (Fig 5E) leading to a significantly reduction of neuronal loss in the gray matter and ventral horn and more intact axons at day 40 post immunization (Fig 5F).

Thus, induction of neuronal mitochondrial activity represents an attractive neuroprotective strategy in CNS inflammation by compensating mitochondrial dysfunction.

## DISCUSSION

Here we show that neuronal oxidative phosphorylation is compromised during CNS inflammation which directly contributes to neuronal vulnerability and can be counteracted by induction of neuronal mitochondrial activity.

In an unsupervised survey we discovered that expression of mitochondrial genes is massively suppressed in motor neurons during CNS inflammation (Schattling *et al*, 2019) and here in particular genes that participate in oxidative phosphorylation. This sparked our interest to explore upstream mechanisms that coordinate this mitochondrial shutdown and whether counter-regulation could rescue neuronal integrity. Our observation is in concordance with other reports that showed alterations of mitochondria in the brain of MS patients with compromised oxidative phosphorylation in demyelinated axons and neurons (Mahad *et al*, 2009; Campbell *et al*, 2011). This could potentially contribute to mitochondrial swelling and dysfunction of axons in EAE (Nikić *et al*, 2011; Sadeghian *et al*, 2016). While we could not detect a decrease in somatic neuronal mitochondrion numbers in the spinal cord during EAE, we discovered a compromised complex IV activity in the spinal cord gray matter. This compromised oxidative phosphorylation during CNS inflammation is likely coordinated by inactivation of PGC-1α, a transcriptional coactivator that acts as a master switch of mitochondrial function (Puigserver *et al*, 1998; Mootha *et al*, 2003). However, in contrast to MS patients, in which a marked decrease in cortical PGC-1α expression has been reported (Witte *et al*, 2013), we could not detect a regulation of PGC-1α in inflamed mouse motor neurons.

By contrast, our data show that CNS inflammation results in a posttranslational modification of PGC-1α (Fernandez-Marcos & Auwerx, 2011). Increased phosphorylation of serine 570 leads to an inactivation of PGC-1α, which could be mediated by increased AKT activity (Li *et al*, 2007). Three different isoforms of AKT exist. Whereas AKT1 and AKT3 are mainly expressed in neurons, AKT2 is predominantly expressed in astrocytes (Levenga *et al*, 2017). So far, the exact function of AKT in neurons is not clear, but AKT1 has been implicated in late long term potentiation (Levenga *et al*, 2017) and AKT3 in neuronal growth (Rivière *et al*, 2012; Adams *et al*, 2016). AKT is activated by phosphatidylinositol 3-kinase (PI3K) and the PI3K/AKT signaling pathway is known to reduce apoptosis and promote survival and proliferation (Brunet *et al*, 1999) and thereby counteracts neuronal cell death (Peltier *et al*, 2007). As TNF-α activates the PI3K/AKT pathway (Osawa *et al*, 2001; Li *et al*, 2017; Gu *et al*, 2006), upregulation of AKT could function as an immediate strategy of neurons against cell death during inflammation. As a trade-off, prolonged downregulation of mitochondrial gene transcripts will reduce ATP levels that could limit neuronal integrity during chronic inflammation. Thus, tuning *Pgc-1α* could act as a switch that modulates oxidative phosphorylation by regulating respective mitochondrial genes under challenging environmental conditions, such as CNS inflammation.

This seems to be particularly relevant in tissues with a high energy demand, e.g. muscle, liver and CNS in which *Pgc-1α* is highly expressed. While astrocytes are mostly relying on glycolysis, neurons mainly generate their energy by oxidative phosphorylation with low glycolytic capacities (Camandola & Mattson, 2017). Therefore, mitochondrial damage and inactivation is particular deleterious for neuronal metabolism. That *Pgc-1α* is important for neuronal health is supported by our observation of an aggravated EAE disease course in neuron-specific *Pgc-1α*-deficient mice. This is in accordance with the global *Pgc-1α* knockout mice that showed neurological symptoms, such as myoclonus, dystonia and limb clasping that was attributed to axonal striatal degeneration (Lin *et al*, 2004), although we were not able to detect this phenotype in our neuron-specific *Pgc-1α* knockout mice. By contrast, our data show that neuronal overexpression of *Pgc-1α* elevated mitochondrial biogenesis and an improved activity of neuronal oxidative phosphorylation, particularly complex IV, which also resulted in a higher overall mitochondrial membrane potential. Notably, these alterations equipped mitochondria with an improved calcium buffering capacity. Besides MS, rise in neuronal intracellular calcium is one of the hallmarks of neurodegenerative diseases (Mattson, 2007). Several mechanisms are relevant that lead to intracellular neuronal Ca^2+^ accumulation in EAE (Siffrin *et al*, 2015), among them upregulation of voltage-gated sodium channels, Na^+^-Ca^2+^ antiporters (Craner *et al*, 2004), Ca^2+^ permeable ion channels (Friese *et al*, 2007; Schattling *et al*, 2012) and nanoscale ruptures of the axonal plasma membrane (Witte *et al*, 2019). Therefore, the improved calcium buffering capacity in *Pgc-1α*^T/T^ neurons could efficiently rescue calcium overload and neuronal demise, which was indicated by an increased neuronal survival during EAE. Calcium uptake by mitochondria holds the promise as a therapeutic strategy for several neurodegenerative diseases (Lee *et al*, 2018; Parone *et al*, 2013). As neuronal mitochondrial deficiency is also associated with other neurodegenerative diseases, e.g. Alzheimer’s disease (Pannaccione *et al*, 2020), induction of mitochondrial activity could serve as a unifying neuroprotective approach (Murphy & Hartley, 2018). Similarly, downstream targets of *Pgc-1α* are repressed in dopaminergic neurons of Parkinson's disease patients (Zheng *et al*, 2010) in which neuronal overexpression of *Pgc-1α* protects dopaminergic neurons in its mouse model (Mudò *et al*, 2012).

As we detected a higher neuronal resilience in *Pgc-1α*^T/T^ mice during CNS inflammation, pharmacological increase of neuronal mitochondria and oxidative phosphorylation could be a promising neuroprotective strategy. Some drugs have already been described to induce *Pgc-1α* in different tissues (Ye *et al*, 2012; Hofer *et al*, 2014; Dumont *et al*, 2012; Noe *et al*, 2013), however we could not confirm neuronal induction of *Pgc-1α* by treating mice with bezafibrate or using transient receptor potential cation channel subfamily V member 4 (*Trpv4*) knockout mice (data not shown). Another possible drug candidate is sirtuin-1 that activates PGC-1α by deacetylation and its neuronal overexpression leads to an ameliorated EAE disease course (Nimmagadda *et al*, 2013). Besides extrinsic induction or activation of *Pgc-1α*, it can also be induced by intrinsic mechanisms, such as cold exposure or exercise (Lin *et al*, 2005). Exercise ameliorates the EAE disease course (Klaren *et al*, 2014; Rossi *et al*, 2009) and shows several benefits in MS patients (Motl *et al*, 2017), however the proof of a neuroprotective effect that is mediated by *Pgc-1α* induction is currently outstanding.

Taken together, we provide evidence for a therapeutic potential of inducing mitochondrial activity in inflammation-induced neurodegeneration, supporting further studies that aim at finding drugs to target this pathway in neurons.

## MATERIAL AND METHODS

### Mice

All mice (C57BL/6J wild type (The Jackson Laboratory), Thy1-Flag-*Pgc-1α* (*Pgc-1α^T^*^/T^) mice on a C57BL/6J genetic background provided by D. Lindholm (Mudò *et al*, 2012), Eno2^Cre+^ mice (The Jackson Laboratory), *Pgc-1α*^flx/flx^ mice (The Jackson Laboratory), Chat-L10a-eGFP mice (Heiman *et al*, 2008) were kept under specific pathogen-free conditions in the central animal facility of the University Medical Center Hamburg-Eppendorf, Hamburg, Germany. Neuron-specific knockout mice were generated by crossing *Pgc-1α^flx^*^/flx^ with Eno2^Cre+^ mice. Animals were housed in a facility with 55–65% humidity at 24 ± 2 °C with a 12-hour light/dark cycle and had free access to food and water. Sex- and age-matched adult animals (10–20 weeks of age) were used in all experiments.

### EAE induction

Mice were anaesthetized with isoflurane 1% to 2% v/v oxygen and immunized subcutaneously with 200 μg myelin oligodendrocyte glycoprotein 35–55 (MOG_35–55_) peptide (peptides & elephants) in complete Freund’s adjuvant (BD) containing 4 mg ml^−1^ Mycobacterium tuberculosis (BD). 200 ng pertussis toxin (Merck) was injected intravenously on the day of immunization and 48 hours later. Animals were scored daily for clinical signs by the following system: 0, no clinical deficits; 1, tail weakness; 2, hind limb paresis; 3, partial hind limb paralysis; 3.5, full hind limb paralysis; 4, full hind limb paralysis and forelimb paresis; 5, premorbid or dead. Animals reaching a clinical score ≥ 4 were sacrificed according to the regulations of the Animal Welfare Act. The experimenters were blinded to the genotype until the end of the experiment, including data analysis. Sex- and age-matched adult animals (8–12 weeks of age) were used in all experiments. For *Pgc-1α^T^*^/T^ mice and wild type controls, three independent EAE experiments were conducted, the data were pooled for final analysis. For *Pgc-1α^flx^*^/flx^ × Eno2^Cre+^ EAE and *Pgc-1α^flx^*^/flx^ controls one EAE was conducted. For analysis of the disease course and weight we only included animals which received a score ± 1 until day 15 and survived until the end of the experiment. Animals were either analyzed at acute stage of EAE (day 12 to day 16 after immunization) or chronic stage (day 30 to day 40 after immunization).

### Gene set enrichment analysis

Gene signature of motor neurons during CNS inflammation was generated by ranking all expressed genes by the DESeq2-derived t statistics based on the comparison of healthy versus EAE motor neurons (Schattling *et al*, 2019). Enrichment analysis was performed using the function ‘gseGO’ of the R package clusterProfiler (Yu *et al*, 2012) on biological process gene ontology terms with at least 50 members. Gene sets with a Benjamini-Hochberg adjusted P value < 0.05 were considered significant and ordered by their normalized enrichment score, with positive value indicating enrichment and negative values indicating de-enrichment. Core enrichment genes driving the de-enrichment of the term ‘GO:0022900 electron transport chain’ were extracted from clusterProfiler results and plotted as heatmap of gene expression values after variance stabilizing transformation. Plotting was performed with the R packages ggplot2, clusterProfiler and tidyheatmap.

### Pgc-1α target identification

For the identification of *Pgc-1α-*induced downstream targets a published microarray dataset of *Pgc-1α*-overexpressing SH-SY5Y cells was used (Lucas *et al*, 2014). The expression matrix was downloaded via GEOquery (GSE100341) and analyzed using limma. Genes with a FDR-adjusted P value < 0.05 and a log2 fold change > 1 in *Pgc-1α* overexpression were considered positively regulated *Pgc-1α* targets.

### Mouse tissue preparation and histopathology of mice

Mice were anaesthetized intraperitoneally with 100 μl solution (10 mg ml^−1^ esketamine hydrochloride (Pfizer), 1.6 mg ml^−1^ xylazine hydrochloride (Bayer) dissolved in water) per 10 g of body weight. For histopathology and immunohistochemistry mice were perfused with 4% paraformaldehyde (PFA), spinal cord tissue was dissected, fixed for 30 min with 4% PFA and then transferred to 30% sucrose in PBS at 4°C. We transversely cut midcervical spinal cord sections at 12 μm with a freezing microtome (Leica Jung CM3000) and stored them in cryoprotective medium (Jung) at –80 °C. For transmission electron microscopy mice were perfused with 4% paraformaldehyde (PFA), 1% glutaraldehyde (GA) in 0.1 M phosphate buffer, spinal cords were dissected and fixed with 2% PFA, 2.5% GA in 0.1 M phosphate buffer. Antibodies directed against CD3 (rabbit IgG, Abcam; RRID: AB_443425), Mac-3 (rat IgG, BD Biosciences; RRID: AB_394780), Iba1 (rabbit, WAKO;, RRID: AB_839504) were visualized by avidin-biotin technique with 3,3-diaminobenzidine (DAB, Sigma) according to standard procedures of the UKE Mouse Pathology Facility using the ultraView Universal DAB Detection Kit (Ventana). For complex IV/cytochrome c oxidase (COX) histochemistry, sections were incubated for 60 min at 37 °C with COX reaction media (diaminobenzidine tetrahydrochloride, cytochrome c and bovine catalase (all from Sigma) in 0.2 M phosphate buffer) and embedded with Aquatex (Merck). Images of tissue sections at ×200 magnification were scanned using Zeiss MIRAX MIDI Slide Scanner (Carl Zeiss, MicroImaging GmbH, Germany). We quantified numbers of inflammatory foci per spinal cord cross-section on hematoxylin and eosin–stained sections. CD3^+^, IBA1^+^ and MAC-3^+^ infiltrating cells were analyzed with ImageJ software (NIH), with the same settings across all experimental groups. For quantification of complex IV activity, background was subtracted and mean gray value intensities of COX reactivity in complete gray matter or the ventral horn were measured by Image J. COX reactivity was normalized to corresponding neuronal count.

### Immunohistochemistry

Sections were incubated in blocking solution (10% normal donkey serum in PBS) containing 0.05% Triton X-100 at room temperature for 45 min and subsequently stained them overnight at 4 °C with antibodies against the following structures: phosphorylated neurofilaments (SMI 31, mouse, 1:500; Covance; RRID: AB_10122491), non-phosphorylated neurofilaments (SMI 32, mouse, 1:500; Covance; RRID: AB_2564642), neuronal nuclei (chicken 1:500; Millipore; RRID: AB_11205760), PGC-1α (PGC-1α, rabbit, 1:100; Novus Biologicals; RRID: AB_1522118) and FLAG sequence (FLAG, mouse, 1:200; Sigma; RRID: AB_259529). As secondary antibodies we used Alexa Fluor 488–coupled donkey antibodies recognizing chicken IgG (1:800, Jackson; RRID: AB_2340375, Cy3-coupled donkey antibodies recognizing mouse IgG (1:800, Jackson; RRID: AB_2340816) and Alexa Fluor 647–coupled donkey antibodies recognizing rabbit IgG (1:800, Abcam; RRID: AB_2752244). We analyzed the sections with a Zeiss LSM 700 confocal microscope. For quantification of neurons and axons tile scans of each animal were taken. Numbers of neurons were manually counted in two defined areas (250 μm × 250 μm) from the central gray matter or ventral horn. For absolute numbers of neurons in the ventral horn, all neurons in both ventral horns per animals were counted. Axons were quantified in three defined areas (90 μm × 90 μm) with ImageJ software using fixed threshold intensity across experimental groups for each type of tissue examined.

### Immunofluorescence

Cortical neurons (DIV 14) were fixed with 4% PFA for 30 min at room temperature, permeabilized with 0.05% Triton and blocked with 10% normal donkey serum in PBS. Cells were stained overnight at 4 °C with antibodies directed against PGC-1α (PGC-1α, rabbit, 1:200; Novus Biologicals; RRID: AB_1522118) and microtubule-associated protein 2 (MAP2, chicken, 1:2500; Abcam; RRID: AB_2138153). As secondary antibodies we used Alexa Fluor 488–coupled donkey antibodies recognizing chicken IgG (1:800, Jackson; RRID: AB_2340375, Cy3-coupled donkey antibodies recognizing mouse IgG (1:800, Jackson; RRID: AB_2340816) and Alexa Fluor 647–coupled donkey antibodies recognizing rabbit IgG (1:800, Abcam; RRID: AB_2752244). Images were taken with a Zeiss LSM 700 confocal microscope.

### Transmission electron microscopy

After dissection and fixation, spinal cords were cut in 1 mm transverse sections and post-fixed in 1% osmium tetroxide (OsO_4_) for 1 hour. Hereafter, samples were dehydrated in serial dilutions of alcohol and embedded with Epon (LX‐112 Resin, Ladd research, USA). Next, ultrathin (50‐85 nm) sections were collected on 200mesh copper grids and counterstained with uranyl acetate and lead citrate. Images of individual motor neurons were taken at ×5000 magnification using a JEM 1010 transmission electron microscope (JEM 1010, Jeol USA). From each motor neuron, we quantified the surface area of neuronal cytoplasm and all cytoplasmic mitochondria using ImageJ (NIH), which were used to calculate mitochondrial density per neuron.

### Primary neuronal culture

Primary hippocampal and cortical cultures were prepared from E16.5 embryos. Hippocampus and cortex were harvested, cut into smaller pieces and incubated in 0.05% Trypsin-EDTA (Gibco) for 6 min at 37 °C. Trypsination was stopped by DMEM-F12 containing 10% FCS. Afterwards, tissue was dissociated in HBSS and centrifuged for 2 min at 500 × g. The pellet was resuspended in Primary Growth Medium (PGM) and cells were plated at 5 × 10^4^ per cm^2^ (hippocampal neurons) or 7.5 × 10^4^ per cm^2^ (cortical neurons) on poly-d-lysine-coated cell culture plates. We maintained cells in PNGM (Primary Neuron Growth Medium BulletKit, Lonza) or neurobasal plus medium (supplemented with B27, penicillin, streptomycin and L-glutamine; Gibco) at 37 °C, 5% CO_2_ and a relative humidity of 98%. To inhibit glial proliferation, we added cytarabine (Sigma, 5 μM) or floxuridine/uridine (Sigma, 10 μM) at 3-4 days in vitro (DIV) at 5 μM and maintained cultures for 14–21 days in vitro (DIV 14 or DIV 21).

### Calcium imaging

Cortical primary neurons were transduced with genetically encoded cytosolic calcium indicator GCaMP6f via adeno-associated virus (AAV7) at day 7 in vitro. Alternatively, chemical calcium indicator Fluo4-AM (ThermoFisher, F14201) was used according to the manufacturer’s instructions (Yasuda *et al*, 2004). Calcium imaging was performed from cortical primary neuronal culture from E16.5 embryos (DIV 15). For details see Appendix Supplementary Methods.

### Live-cell metabolic assay

Hippocampal neurons were seeded in a poly-d-lysine-coated XF 96-well cell culture microplate (Seahorse Bioscience, Copenhagen, Denmark) in triplicate at 5 × 10^5^ cells per well in 1 ml neuronal growth medium and then incubated at 37 °C in 5% CO_2_. At DIV14, media from neurons was removed, replaced by 180 μl of assay media (Assay Media from Bioscience with 25 mM glucose, 1 mM sodium pyruvate and 2 mM L-glutamine; pH 7.4) and incubated in a CO_2_ free incubator at 37 °C for 1 hour. Compounds (2 μM oligomycin, 1 μM FCCP, 0.5 μM rotenone/antimycin A, all in assay media) were added into the appropriate ports of a hydrated sensor cartridge. For baseline measurement control ports were left without compound addition. Cell plate and cartridge were then placed into the XFe96 Analyzer and results analyzed by WAVE Software.

### Cell viability assay (calcein)

Hippocampal neurons were seeded in a poly-d-lysine-coated XF 96-well cell culture microplate (Seahorse Bioscience, Copenhagen, Denmark) in quadruplicate at 5 × 10^5^ cells per well in 1 ml neuronal growth medium and then incubated at 37 °C in 5% CO_2_. At DIV14 calcein-AM (Sigma) was added to the cultures (2 μM), incubated at 37 °C in 5% CO_2_ for 30 min, washed in optic buffer (ddH_2_O supplemented with 10 mM glucose, 140 mM NaCl, 1 mM MgCl_2_, 5 mM KCl, 20 mM HEPES, 2 mM CaCl_2_, pH 7.3 adjusted with NaOH) at room temperature and fluorescence intensity was measured according to manufacturer’s instructions.

### Mitochondrial membrane potential assay

Primary neurons were seeded on μ-dishes (ibidi) and tetramethylrhodamin-ethylester (TMRE) assay (Abcam) was performed on DIV14 according to manufacturer’s protocol. Cells were treated with 10 nM TMRE for 20 min in PGM, washed twice with HBSS w/o phenol red (Gibco) and then equilibrated for 10 min in HBSS w/o phenol red and live imaged at 555 nm with a Zeiss LSM 700 confocal microscope. For internal control, 20 μM FCCP was added 10 min prior to TMRE. Images were taken with a Zeiss LSM 700 confocal microscope. Total fluorescence of five cells per biological replicate were analyzed.

### Translating ribosome affinity purification (TRAP)

TRAP was performed as previously described (Schattling *et al*, 2019). For details see Appendix Supplementary Methods.

### Fluorescence-activated cell sorting

CD4+ T cells (CD45+CD3+CD4+), CD8+ T cells (CD45+CD3+CD8+), B cells (CD45+CD19+), macrophages (CD45+CD3-CD19-Ly6G-NK1.1-CD11b+ F4/80+) and microglia (CD45med CD11b+were sorted into collection tubes coated with FCS and filled with complete RPMI 1640 medium (1% penicillin, streptomycin, 0.1% 2-ME) with 20% FCS using a FACSAria device (BD Biosciences). Purity of sorted populations was routinely above 95%. Then cells were pelleted by centrifugation at 300 × g for 10 min at 4°C, dry frozen in liquid nitrogen, and stored at –80°C until RNA isolation. For details see Appendix Supplementary Methods.

### Quantitative real-time PCR

RNA was purified using the RNeasy Mini Kit (Qiagen) and reverse transcribed to cDNA with the RevertAid H Minus First Strand cDNA Synthesis Kit (Thermo Fisher) according to the manufacturer’s instructions. Gene expression was analyzed by quantitative real-time PCR performed in an ABI Prism 7900 HT Fast Real-Time PCR System (Applied Biosystems) using TaqMan Gene Expression Assays (Thermo Fisher) for *Ppargc1a* (*Pgc-1α*; Mm00464452_m1), *Alas* (Mm01235914_m1), *Cox4i1* (Mm01250094_m1), *Cs* (Mm00466043_m1), *Nrf1* (Mm01135606_m1), *Tfam* (Mm00447485_m1) and *Tbp* (Mm01277042_m1). Gene expression was calculated as 2^−ΔCT^ relative to *Tbp* as endogenous control.

### mtDNA copy number quantification

DNA was purified using the DNeasy Blood and Tissue Kit (Qiagen) according to the manufacturer’s instructions. To destroy RNA, samples were treated with RNAse (Qiagen). Mitochondrial DNA copy numbers relative to diploid chromosomal DNA content was analyzed by quantitative real-time PCR performed in an ABI Prism 7900 HT Fast Real-Time PCR System (Applied Biosystems) using TaqMan Gene Expression Assays (Thermo Fisher) for *Cox2I* (Mm03294838_g1) and *β-actin* (Mm_00607939_s1). MtDNA copy numbers were quantified as 2^−ΔCT^ relative to β-actin.

### Pgc-1α DNA copy number quantification

DNA was lysed with 50 μl QuickExtract™ DNA Extraction Solution (Lucigen). Copy numbers were analyzed by quantitative real-time PCR performed in an ABI Prism 7900 HT Fast Real-Time PCR System (Applied Biosystems) using TaqMan Copy Number Assays (Thermo Fisher) for *Ppargc1a* (*Pgc-1α,* Mm00164544_cn, FAM) and *Tfrc* (TaqMan™ Copy Number reference Assay for mouse, VIC) in a duplex PCR. *Pgc-1α* copy numbers were quantified as 2^−ΔCT^ relative to *Tfrc*.

### Immunoblot

Denatured spinal cord samples were loaded onto a Bolt™ 4–12% Bis-Tris Plus Gel (Invitrogen™) in a Mini Gel Tank (Life Technologies) and proteins were separated. For immunodetection the primary antibodies against the following proteins were used: PGC-1α (rabbit 1:2000; Novus Biological; RRID: AB_1522118), phosphorylated PGC-1αS570 (rabbit 1:1000; R&D Systems; RRID: AB_10890391) and β-actin (rabbit 1:1000; Cell Signaling Technology; RRID :AB_330288). For details see Appendix Supplementary Methods.

### Study approval

All animal care and experimental procedures were performed according to institutional guidelines and conformed to requirements of the German Animal Welfare Act. All animal experiments were approved by the local ethics committee (Behörde für Soziales, Familie, Gesundheit und Verbraucherschutz in Hamburg; G22/13 and 122/17). We conducted all procedures in accordance with the ARRIVE guidelines (Kilkenny *et al*, 2010).

### Statistics

Experimental data were analyzed using Prism 8 software (GraphPad) and are presented as mean values ± SEM. Statistical analyses were performed using the appropriate test indicated in the figure legends Shapiro Wilk test was used to analyze normality. Unless stated otherwise, differences between two experimental groups were determined by unpaired, two-tailed Mann-Whitney or Students t-test. Differences between three experimental groups were determined by multiple comparisons test following one-way ANOVA. Significant results are indicated by asterisk: *P < 0.05, **P < 0.01, ***P < 0.001, ****P < 0.0001.

## ACKNOWLEDGEMENTS

We thank the UKE Mouse Pathology Facility for histopathology of EAE mice, the UKE Vector Facility for generating the Calcium indicator viruses and the animal facility. We thank Hilmar Bading and Peter Bengtson for advice on calcium imaging. SCR is supported by a Clinician-Scientist Fellowship from the Stifterverband für die Deutsche Wissenschaft and the Hertie Network of Excellence in Clinical Neuroscience of the Gemeinnützige Hertie-Stiftung. This work is supported by grants of the Deutsche Forschungsgemeinschaft (DFG) to MAF an MF (FR1720/9-1, FR1720/9-2, FR1638/3-1, FR1638/3-2) which are part of the DFG FOR2289.

## AUTHOR CONTRIBUTIONS

SCR and MAF designed the experiments for the study and wrote the manuscript. SCR, AS and MW analyzed the data. SCR, AS, ST, VV, VR, NP, SB, CS, LB and BS performed experiments. JBE conducted bioinformatics analysis. SCR, AS, MW, OP, MF, MAF interpreted the data. MAF conceived and supervised the study.

## EXPANDED VIEW FIGURE LEGENDS

**Figure EV1.**

(**A**) Representative transmission electron microscopy (TEM) images of cervical spinal cord motor neurons of healthy and chronic EAE mice. Scale bar: 5 μm.

(**B**) Quantification of transmission electron microscopy (TEM) analysis of motor neuronal mitochondrial numbers of healthy (n = 3) and chronic EAE (n = 3) mice (2–3 cells per mice). Bars show mean values ± s.e.m.

(**C**) Quantification of transmission electron microscopy (TEM) analysis of motor neuronal mitochondrial size of healthy (n = 3) and chronic EAE (n = 3) mice (2–3 cells per mice). Bars show mean values ± s.e.m. Statistical analysis in (**B**) and (**C**) was performed by unpaired, two-tailed Student’s t-test.

**Figure EV2.**

Representative immunoblots of phosphorylated PGC-1αS570 (pPGC-1α) of cervical spinal cords of healthy wild-type mice with and without prior treatment of lambda protein phosphatase for validation of the pPGC-1αS570 antibody.

**Figure EV3.**

(**A**) Relative qPCR mRNA expression of Thy-1 to *Tbp* in hippocampal primary neuronal culture (hPNC) and cortical primary neuronal cultures (cPNC) (DIV14) of wild-type (n = 6 per group) mice. Bars show mean values ± s.e.m.

(**B**) Relative diploid nuclear chromosomal DNA copy numbers of *Pgc-1α* in tail biopsy of wild-type (n = 9) and Pgc-1α^T/T^ (n = 16) mice. Bars show mean values ± s.e.m.

(**C**) Relative qPCR mRNA expression of *Pgc-1α* in hippocampus (Hip), cortex and cervical spinal cord (SC) of wild-type (n = 4) and *Pgc-1α*^T/T^ (n = 4) mice. Bars show mean values ± s.e.m.

(**D**) Relative qPCR mRNA expression of Pgc-1α in hippocampal primary neuronal cultures (hPNC) and cortical primary neuronal cultures (cPNC) (DIV14) of wild-type (n = 3) and *Pgc-1α*^T/T^ (n = 3) mice. Bars show mean values ± s.e.m.

(**E**) Relative qPCR mRNA expression of *Pgc-1α* target genes in hippocampal primary neuronal cultures (hPNC) (DIV14) of wild-type (n = 3) and *Pgc-1α*^T/T^ (n = 3) mice. Bars show mean values ± s.e.m.

(**F**) Relative qPCR mRNA expression of *Pgc-1α* target genes in cortical primary neuronal cultures (cPNC) (DIV14) of wild-type (n = 3) and *Pgc-1α*^T/T^ (n = 3) mice. Bars show mean values ± s.e.m.

(**G**) Relative qPCR mRNA expression of *Pgc-1α* target genes in hippocampus of wild-type (n = 4) and *Pgc-1α*^T/T^ (n = 4) mice. Bars show mean values ± s.e.m

(**H**) Relative qPCR mRNA expression of *Pgc-1α* target genes in cortex of wild-type (n = 4) and *Pgc-1α*^T/T^ (n = 4) mice. Bars show mean values ± s.e.m.

(**I**) Relative qPCR mRNA expression of *Pgc-1α* target genes in cervical spinal cord of wild-type (n = 4) and *Pgc-1α*^T/T^ (n = 4) mice. Bars show mean values ± s.e.m.

(**J**) Representative immunocytochemical staining of PGC-1α in cortical primary neuronal culture (cPNC) (DIV14) of wild-type and *Pgc-1α*^T/T^ mice. Co-stainings for microtubule-associated protein 2 (Map2) and 4′,6-diamidino-2-phenylindole (DAPI). Scale bar: 10 μm.

(**K**) Representative immunohistochemical staining of PGC-1α in spinal cord of wild-type and *Pgc-1α*^T/T^ mice. Co-stainings for neuronal nuclei (NeuN), FLAG and 4′,6-diamidino-2-phenylindole (DAPI). Scale bar: 20 μm.

(**L**) Mean fluorescence intensity (MFI) quantification of calcein of hippocampal primary neuronal culture (hPNC) (DIV14) of wild-type (n = 1) and *Pgc-1α*^T/T^ (n = 1) mice (3 wells per culture) to determine cell viability. Bars show mean values ± s.e.m.

Statistical analysis in (**A**) and (**B**) was performed by unpaired, one-tailed Student’s t-test, in (**C**), (**D**), (**E**), (**F**), (**G**), (**H**), (**I**) and (**L**) by unpaired, two-tailed Student’s t-test. *P < 0.05, **P < 0.01, ***P < 0.001, ****P < 0.0001.

**Figure EV4.**

(**A**) Relative qPCR mRNA expression of *Eno2* to *Tbp* in hippocampal primary neuronal cultures (hPNC) (DIV14) of wild-type (n = 3) mice. Bars show mean values ± s.e.m.

(**B**) Relative qPCR mRNA expression of *Pgc-1α* in sorted immune cells in relation to hippocampal primary neuronal cultures (hPNC) (DIV14) of wild-type (n = 3) mice. Bars show mean values ± s.e.m.

(**C**) Relative qPCR mRNA expression of *Pgc-1α* in hippocampus (Hip), cortex and cervical spinal cord (SC) of *Pgc-1α*^flx/flx^ (n = 3) and *Pgc-1α*^flx/flx^ × Eno2^cre+^ (n = 3) mice. Bars show mean values ± s.e.m.

(**D**) Relative qPCR mRNA expression of *Pgc-1α* target genes in hippocampus of *Pgc-1α*^flx/flx^ (n = 3) and *Pgc-1α*^flx/flx^ × Eno2^cre+^ (n = 3) mice. Bars show mean values ± s.e.m.

(**E**) Relative qPCR mRNA expression of *Pgc-1α* target genes in cortex of *Pgc-1α*^flx/flx^ (n = 3) and *Pgc-1α*^flx/flx^ Eno2^cre+^ (n = 3) mice. Bars show mean values ± s.e.m.

(**F**) Relative qPCR mRNA expression of *Pgc-1α* target genes in cervical spinal cord of *Pgc-1α*^flx/flx^ (n = 3) and *Pgc-1α*^flx/fl^ × Eno2^cre+^ (n = 3) mice. Bars show mean values ± s.e.m

Statistical analysis in (**B**) was performed by multiple comparisons test following one-way ANOVA, in (C), (D), (**E**) and (**F**) by unpaired, two-tailed Student’s t-test. *P < 0.05, **P < 0.01, *** P < 0.001, ****P < 0.0001.

